# A CRISPR interference screen reveals a role for cell wall teichoic acids in conjugation in *Bacillus subtilis*

**DOI:** 10.1101/2021.05.04.442501

**Authors:** M. Michael Harden, Alan D. Grossman

## Abstract

Conjugative elements are widespread in bacteria and include plasmids and integrative and conjugative elements (ICEs). They transfer from donor to recipient cells via an element-encoded type IV secretion system. These elements interact with and utilize host functions for their lifecycles. We sought to identify essential host genes involved in the lifecycle of the integrative and conjugative element ICE*Bs1* of *Bacillus subtilis*. We constructed a library of strains for inducible knockdown of essential *B. subtilis* genes using CRISPR interference. Each strain expressed one guide RNA in ICE*Bs1*. We induced partial interference of essential genes and identified those that caused an acute defect in acquisition of ICE*Bs1* by recipient cells. This screen revealed that reducing expression of genes needed for synthesis of cell wall teichoic acids caused a decrease in conjugation. Using three different ways to reduce their synthesis, we found that wall teichoic acids were necessary in both donors and recipients for efficient conjugative transfer of ICE*Bs1*. Further, we found that depletion of wall teichoic acids caused cells involved in ICE*Bs1* conjugation to die, most likely from damage to the cell envelope. Our results indicate that wall teichoic acids help protect against envelope stress caused by an active conjugation machinery.

## Introduction

Horizontal gene transfer is fundamental to bacterial evolution, allowing for the rapid spread of genes involved in processes as diverse as antibiotic resistance, resource utilization, and pathogenesis (Frost *et al*., 2005; Wiedenbeck and Cohan, 2011; Soucy *et al*., 2015). Horizontal gene transfer is often mediated by conjugative elements which encode machinery to transfer a copy of the element from a host to a recipient via direct cell-to-cell contact. Although most studies of conjugative elements have focused on plasmids, integrative and conjugative elements (ICEs) appear to be the most abundant type of conjugative element and have been identified in every bacterial clade (Guglielmini *et al*., 2011).

ICEs normally reside integrated in a host chromosome where they are passively replicated and inherited. While in this inactive state, most ICE genes are not expressed. ICEs can be activated either stochastically or in response to certain conditions (i.e., DNA damage to the host, resource limitation), at which point a site-specific recombinase excises the ICE from the host chromosome to form a circular plasmid (Bañuelos-Vazquez *et al*., 2017). For conjugative DNA transfer, the ICE plasmid DNA is nicked and a single strand is transferred out of the donor and into a recipient cell through the element-encoded type IV secretion system. Once transferred, the ssDNA becomes double stranded and can integrate into the chromosome of the new host, forming a stable transconjugant (Wozniak and Waldor, 2010; Johnson and Grossman, 2015; Delavat *et al*., 2017).

ICE*Bs1* is a relatively small ICE (∼20 kb) (Auchtung *et al*., 2016) found in most isolates of the gram-positive bacterium *Bacillus subtilis*. Virtually all of the ICE*Bs1* genes required for conjugative transfer have homologs or analogs in other conjugative elements (Bhatty *et al*., 2013; Leonetti *et al*., 2015). ICE*Bs1* is normally integrated into *trnS*-*leu2* (a tRNA gene). When integrated, only a few ICE*Bs1* genes are expressed. Gene expression and subsequent excision are induced during the RecA-dependent SOS response, or in the presence of other *B. subtilis* cells lacking a copy of the element (Auchtung *et al*., 2005; Bose *et al*., 2008). ICE*Bs1* can be experimentally activated, typically in 50-90% of cells in a population by overproducing the element-encoded regulatory protein RapI, enabling high frequencies of transfer that make it quite useful for studying conjugation (Auchtung *et al*., 2005; Lee *et al*., 2007).

The life cycles of all mobile genetic elements depend on various host functions. In this study, we sought to identify essential host genes that are necessary for the life cycle of ICE*Bs1*. We were most interested in identifying host functions that affect conjugation, rather than host functions that affect the production of the ICE proteins (e.g., transcription, translation) or the replication of ICE DNA (Lee *et al*., 2010).

We used CRISPR interference (CRISPRi) to block transcription of essential *B. subtilis* genes and identified host factors in an ICE*Bs1* donor that are important for transfer. Several genes were identified in this screen and we chose to focus on those involved in the synthesis of cell wall teichoic acids.

Wall teichoic acids (WTAs) are polyol-phosphate repeats that are attached to the peptidoglycan cell wall of gram-positive bacteria [reviewed in: (Swoboda *et al*., 2010; Brown *et al*., 2013)]. In *B. subtilis* 168, WTAs comprise 45-60 repeats of glycerol 3-phosphate (Pollack and Neuhaus, 1994). These repeats can be modified by WTA-tailoring enzymes, most notably by D-alanylation and glycosylation. In *B. subtilis* 168, WTAs are synthesized by enzymes encoded by the *tag* genes (*tagO, tagAB, tagDEF*, and *tagGH*). TagO catalyzes the first step of WTA biosynthesis (D’Elia *et al*., 2006). TagA catalyzes the second step and is the first committed step (D’Elia *et al*., 2009). WTAs are not strictly required for cell growth. WTA-depleted *B. subtilis* cells are viable but slow-growing, and have significant alterations in cell shape and cell separation (D’Elia *et al*., 2006). Some WTA biosynthesis genes (*tagBDFGH*) are conditionally essential, likely because their deletion results in either the accumulation of toxic intermediates or the sequestration of vital cellular resources (D’Elia *et al*., 2006; D’Elia *et al*., 2009).

WTAs have multiple functions, including regulating peptidoglycan synthesis and turnover, mediating cell-cell and cell-surface adhesion, and regulating autolysin activity (Brown *et al*., 2013). We found that wall teichoic acids are necessary in both ICE*Bs1* donor and recipient cells for efficient transfer of the element. The activity of the ICE*Bs1* conjugation machinery was toxic to cells that were depleted of wall teichoic acids, and these cells appeared to die from damage to the cell envelope caused by the conjugation machinery.

## Results

### A CRISPRi screen for essential host genes involved in conjugation

#### CRISPRi knockdown of essential genes

We sought to identify host genes in an ICE*Bs1* donor that are important for transfer of the element. We used a *B. subtilis* CRISPRi system (Peters *et al*., 2016) to reduce expression of essential genes in *B. subtilis*, and screened for those that caused a defect in conjugation. Briefly, the system comprises a catalytically dead Cas9 nuclease from *Streptococcus pyogenes* (dCas9) under the regulation of the xylose-inducible promoter Pxyl. Constitutively expressed single guide RNAs (sgRNA) contain a 20 nt region corresponding to the target gene of interest. When dCas9 is produced via the addition of xylose, it complexes with the sgRNA and stably binds to the host gene specified by the targeting region of the sgRNA. This interaction sterically blocks transcript elongation, thereby lowering expression of the targeted gene or operon. The library of sgRNAs used in this study (a gift of Peters et al.) collectively targeted a set of 289 proposed essential *B. subtilis* genes (Peters *et al*., 2016), 257 of which were subsequently verified to be essential in a systematic gene knockout analysis (Koo *et al*., 2017).

#### CRISPRi library in ICE*Bs1*

We created a library of donor strains in which ICE*Bs1* could be induced by overproducing its activator protein RapI. All donor strains contained a xylose-inducible Pxyl-*dcas9* integrated into the host chromosome. Each donor strain also had one constitutively expressed sgRNA allele (Pveg-*sgRNA*) integrated into a site in ICE*Bs1* that is not needed for transfer. In this way, we constructed a pooled library of donor strains with each individual donor strain representing a knockdown of one essential *B. subtilis* gene (Fig. 1A).

**Figure 1.**
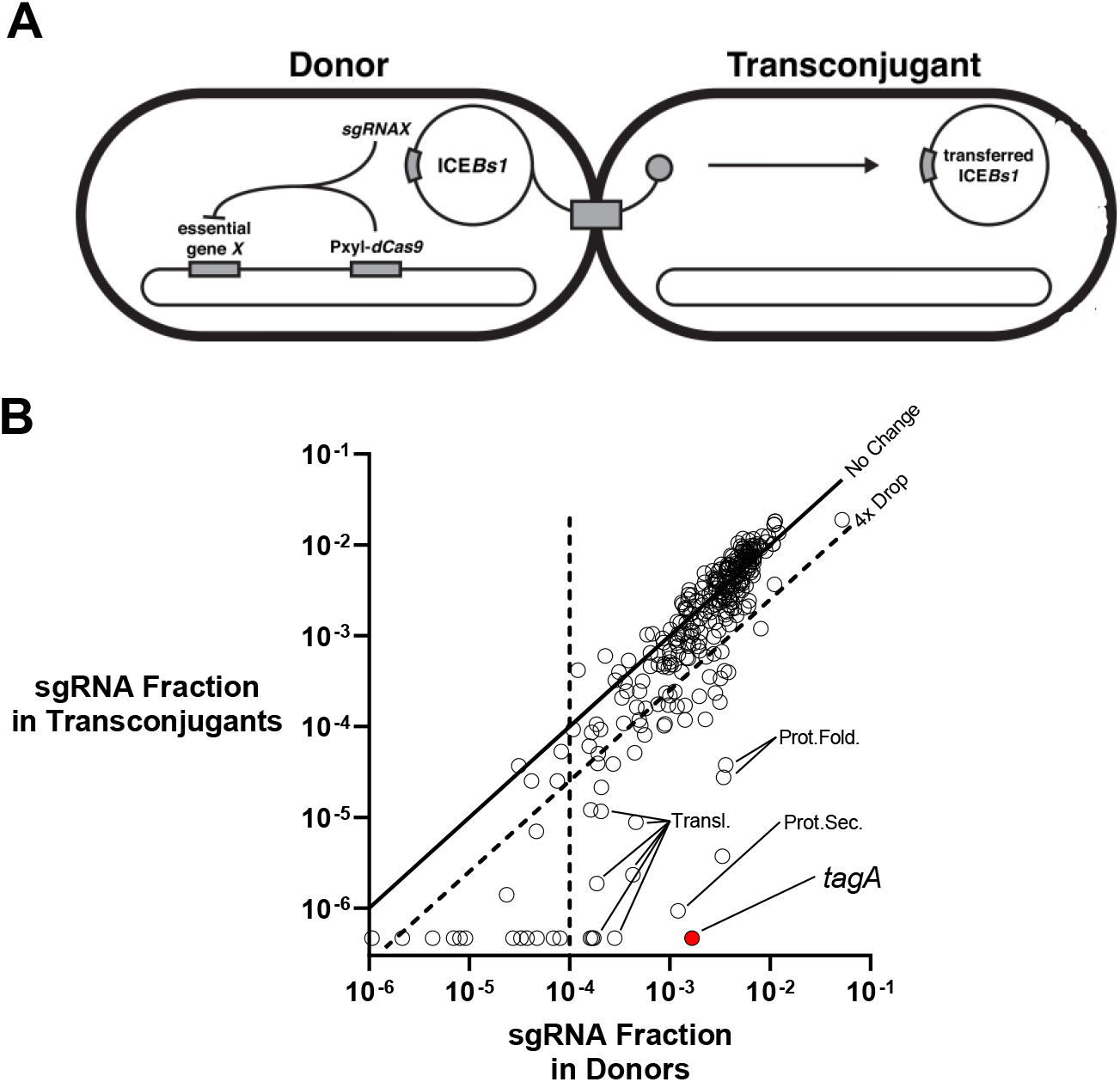
CRISPRi screen identifies essential host gene knockdowns that affect transfer of ICE*Bs1*. **A**. Experimental design of CRISPRi screen. Each ICE*Bs1* donor contains a xylose-inducible Pxyl-*dcas9* allele integrated into a non-ICE locus of the host chromosome and a constitutively expressed sgRNA allele integrated into ICE*Bs1*. Each sgRNA allele has a unique 20 bp targeting region which specifies a knockdown of a single proposed essential gene. A pooled library of inducible ICE*Bs1* donor strains (*lacA*::{P*xyl*-*dcas9* (*ermR*)} Δ*amyE117*::{P*spank*-*rapI* (*spc*)} Δ*rapIphrI*::(*amyE kan*)::{P*veg*-*sgRNA*^*X*^ (*cat*)}), collectively representing partial knockdowns of all *B. subtilis* essential genes, was induced to mate with an ICE*Bs1*^0^ recipient strain (CAL89). If a knockdown results in a transfer defect, the corresponding sgRNA would be less abundant in the transconjugant population relative to the pre-mating donor population. **B**. Results of CRISPRi screen. The pools of sgRNA alleles in the pre-mating donor population and post-mating transconjugant population were sequenced and compared. Each point corresponds to one sgRNA allele included in the screen. The x- and y-axes correspond to the fractions of the donor and transconjugant pools, respectively, that an sgRNA allele represents. The solid diagonal line (x = y) represents no change in abundance between the two populations. Points falling below the diagonal dashed line correspond to alleles that were >4-fold depleted in the transconjugant pool. Points to the left of the vertical dashed line represent sgRNAs that were largely depleted from the donor pool prior to mating (< 0.01%). Select sgRNAs are annotated by name or functional grouping. More detailed results are reported in Table S1.

#### The screen

We used the CRISPRi system in ICE*Bs1* to partially decrease expression of essential genes. We used partial rather than full expression of *dcas9* because we were concerned that high levels, in combination with an sgRNA targeting an essential gene, would cause a significant fitness defect of each donor strain and would have strongly biased or interfered with the screen. Therefore, we grew the library of ICE*Bs1* CRISPRi donors in a low concentration of xylose (0.01%) in rich medium for approximately six generations to cause the partial knockdown of expression of essential genes. We activated ICE*Bs1* in this library by expression of *rapI*, mixed the donors with recipients (strain CAL89), and put the mating mix on a filter for two hours (Experimental Procedures). After recovering cells from the mating filters, we used antibiotic resistance markers in ICE*Bs1* (*kan*) and the recipient chromosome (*str*) to select for transconjugants. Transconjugants from the selection plates were pooled for further analyses. We used next-generation sequencing to determine the relative abundance of sgRNA alleles in a sample of: 1) donors harvested immediately prior to mating and 2) the pool of transconjugants. The overall mating efficiency of this pooled library (0.17%) was similar to that of a control strain with a knockdown targeted to a nonessential gene (MMH233, 0.36%), indicating that the partial knockdown treatment did not have a substantial global effect on conjugation.

Importantly, the target of a gene knockdown is specified by the sgRNA encoded within ICE*Bs1* itself. As a consequence, every transconjugant generated by this library contains a genetic record of the donor strain that produced it. We can determine the relative mating efficiency of a given donor strain by pooling the population of transconjugants, collectively sequencing their sgRNA alleles, and then determining which sgRNAs differ in abundance compared to the pre-mating donor population. If a knockdown compromised ICE*Bs1* mating efficiency, then the sgRNA corresponding to that gene would be underrepresented in the resulting pool of transconjugants. Conversely, if a knockdown improved ICE*Bs1* mating efficiency, then the sgRNA allele corresponding to the that gene would be overrepresented in the pool of transconjugants.

Decreased expression of most of the essential genes had little or no effect on mating efficiency (Fig. 1B). We compared the relative abundance of each sgRNA in the transconjugant population to the pre-mating donor population and found that >80% of knockdowns resulted in less than a 4-fold change in the abundance of the sgRNA gene in the transconjugant pool relative to the starting population (Fig. 1B, points above diagonal dotted line). We did not detect an increase greater than four-fold in any of the sgRNA genes in transconjugants, indicating that none of the essential genes seemed to be substantially inhibiting the function of ICE*Bs1*.

We chose to focus on those genes that caused ≥ 4-fold effects to increase the chances that observed effects were robust and significant. We also focused on knockdown strains that were well represented in the donor pool (Fig. 1B, points to the right of the horizontal line, >0.01%),and that had the largest decrease in conjugation. Of this subset of knockdowns, most were involved in processes that directly affect the production of the conjugation machinery, including genes involved in translation, protein secretion, and protein folding. We did not study these. In contrast, knockdown of *tagA* resulted in the most severe transfer defect out of all genes tested in the screen. Knockdowns of two other WTA biosynthesis genes (*tagD* and *tagF*) also resulted in >4-fold defects in conjugation (∼10-and ∼4.5-fold respectively; Table S1). We decided to further investigate the role of *tagA* and WTA biosynthesis in conjugation.

### WTAs are necessary in an ICE*Bs1* donor for efficient transfer

To validate the apparent effect of *tagA* revealed in the CRISPRi screen, we directly tested for effects of *tagA* on ICE*Bs1* conjugation in several ways. First, we measured the conjugation efficiency {stable transconjugants per donor; each measured as colony-forming units (CFUs)} from a homogenous population of donor cells in which *tagA* expression was inhibited by CRISPRi. This is in contrast to the screen which used a population of strains representing the entire CRISPRi library. As anticipated, decreasing *tagA* expression in an ICE*Bs1* donor resulted in an acute drop in conjugation (Fig. 2A). The severity of the defect increased as *tagA* expression decreased, with the strongest knockdown resulting in no detectable ICE*Bs1* transfer (< 1 x 10^−3^).

**Figure 2.**
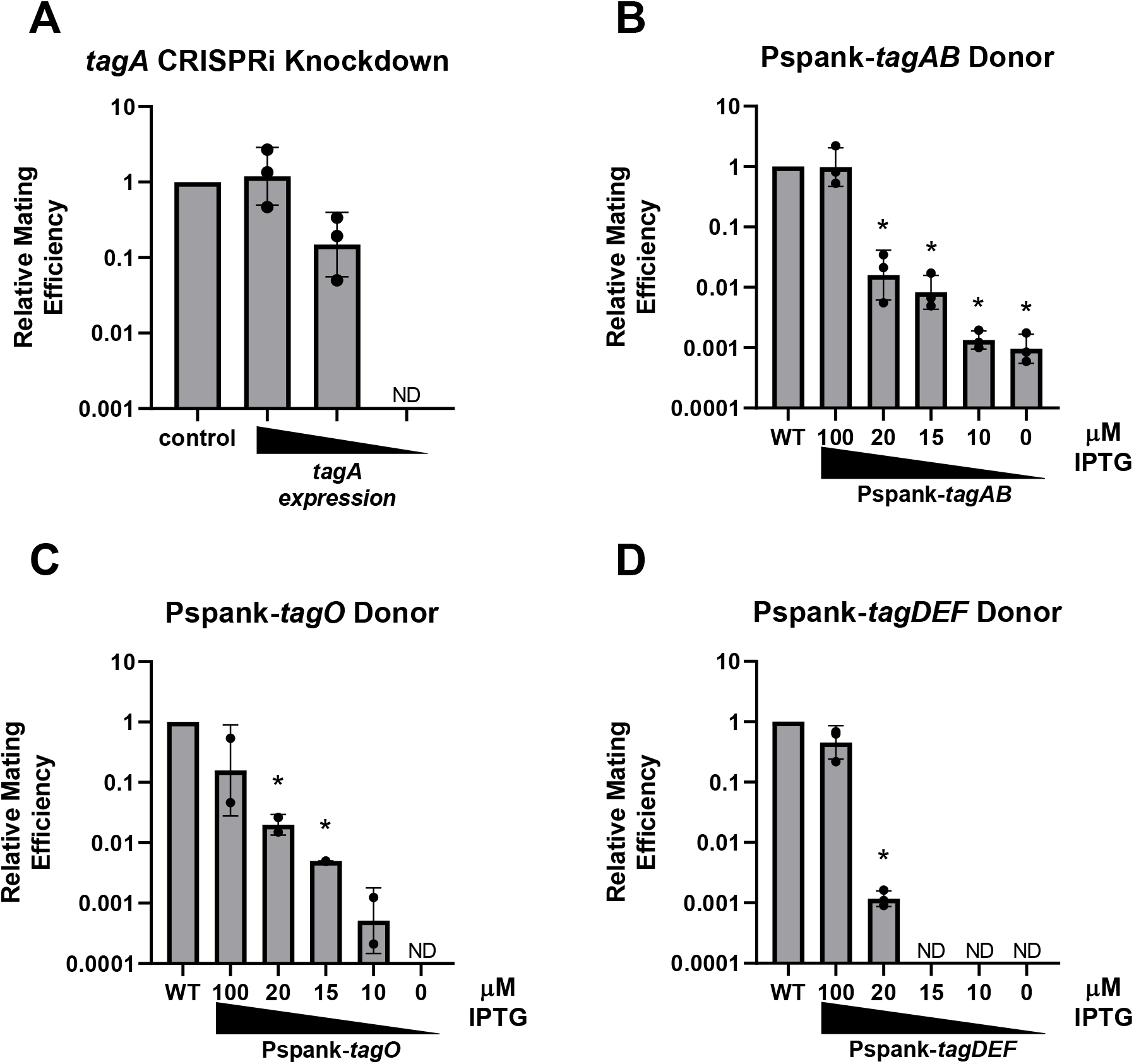
Decreased expression of *tag* genes causes a decrease in conjugation. Expression of *tag* genes was inhibited in ICE*Bs1* donors and the resulting impact on mating efficiency was examined. Relative mating efficiencies are reported as number of transconjugants per pre-mating donor normalized to a same-day control. Graphs show averages and standard deviations of log-transformed data from two or three biological replicates. * indicates average relative mating efficiency was significantly different from 1 (Student’s one-sample t-test, p < 0.05). **A**. An individual ICE*Bs1* donor strain with a xylose-inducible CRISPRi knockdown of *tagA* (MMH527) was induced to mate with an ICE*Bs1*^0^ recipient strain (CAL89). Knockdowns were induced via incubation with either 0, 0.01%, or 0.1% xylose. An ICE*Bs1* donor strain with a control knockdown was used as the control (MMH525, *cgeD*, 0.1% xylose). ND indicates mating efficiency was below limit of detection (< .001). Average mating efficiency of the control was 8.80 x 10^−4^ with a standard deviation of 4.11 x 10^−4^. **B-D**. ICE*Bs1* donor strains were constructed in which a WTA biosynthesis operon, either *tagAB* (B; MMH578), *tagO* (C; MMH577), or *tagDEF* (D; MMH608), was placed under the control of the IPTG-inducible promoter Pspank. Each donor was mated with an ICE*Bs1*^0^ recipient strain (CAL89). Expression of the *tag* operon was controlled by growing the donor strain at the indicated IPTG concentration. A wild type ICE*Bs1* donor (CAL874) was used as a control. ND indicates mating efficiency was below limit of detection (< .0006 for C; < 0.002, 0.002, and 0.001 for the 15 µM, 10 µM, and 0 µM treatments in D, respectively). Average mating efficiency and standard deviation of the controls were 7.78 x 10^−4^ ± 3.96 x 10^−4^, 6.94 x 10^−4^ ± 6.20 x 10^−4^, and 6.58 x 10^−4^ ± 3.66 x 10^−4^ for B, C, and D respectively.

We inhibited WTA synthesis in ICE*Bs1* donors in two additional ways. In one, we made three different donor strains, each with one of three *tag* operons (*tagO, tagAB*, or *tagDEF*) under the control of the LacI-repressible-isopropyl-ß-D-thiogalactopyranoside (IPTG)-inducible promoter Pspank. We grew cells in different IPTG concentrations (ranging from 0 to 100 µM) for approximately six generations and then measured effects on conjugation. Reducing expression of any of the three operons (*tagO, tagAB, tagDEF*) resulted in an acute drop in conjugation (Fig. 2B-D). In all three cases, mating efficiency dropped more than 1,000-fold at the lowest level of expression tested (no IPTG).

We also inhibited WTA synthesis using a low concentration (1 μg/ml) of the antibiotic tunicamycin. At this concentration, tunicamycin specifically inhibits WTA biosynthesis in gram-positive bacteria by blocking the activity of TagO (Pooley and Karamata, 2000; Campbell *et al*., 2011). At higher concentrations (>10 µg/ml) it inhibits peptidoglycan biosynthesis and cell growth (Price and Tsvetanova, 2007; Campbell *et al*., 2011). Importantly, inhibiting WTA biosynthesis with tunicamycin (1 µg/ml) for approximately three generations (60 minutes) in an ICE*Bs1* donor decreased ICE*Bs1* transfer ∼150-fold (Fig. 3A). Based on these results, we infer that WTAs are likely necessary for efficient transfer of ICE*Bs1*. Alternatively, it is formally possible that the biosynthetic activity of TagO and WTA biosynthesis per se is important and not accumulation of WTAs themselves.

**Figure 3.**
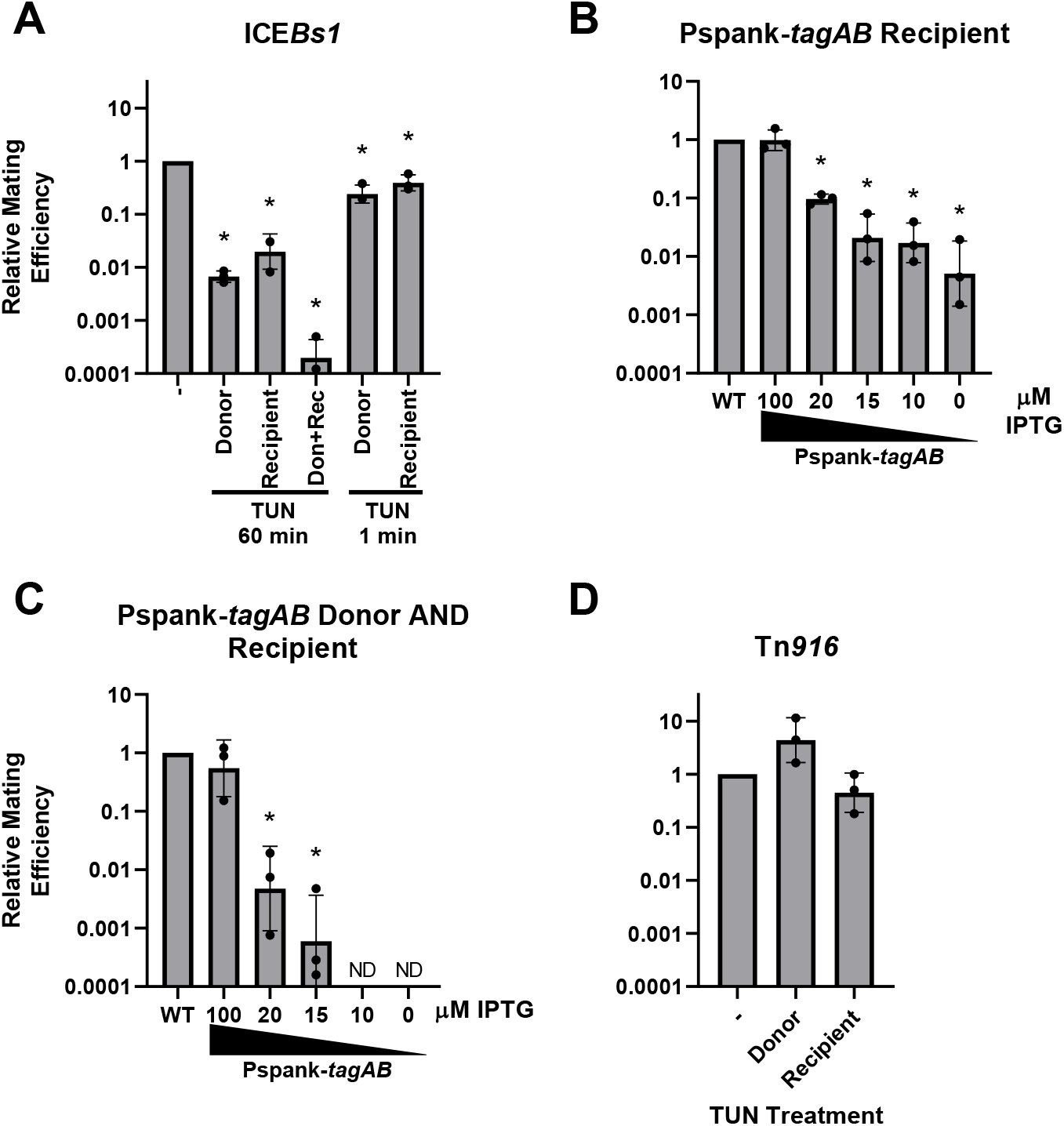
WTAs are necessary in both ICE*Bs1* donors and recipients for efficient transfer of the element. WTA biosynthesis was inhibited in ICE donors and ICE^0^ recipients, and the resulting impact on mating efficiency was examined. Relative mating efficiencies are reported as described in Figure 2. Graphs show averages and standard deviations of log-transformed data from three biological replicates. * indicates average relative mating efficiency was significantly different from 1 (Student’s one-sample t-test, p < 0.05). **A**. WTA biosynthesis was inhibited in an ICE*Bs1* donor (MMH550), an ICE*Bs1*^0^ recipient (MMH676), or both by adding the TagO-inhibiting drug tunicamcyin (1 μg/ml). The drug was added either 60 minutes prior to mating to block synthesis and deplete WTAs, or 1 minute prior to mating to block WTA synthesis without substantial WTA depletion. Average mating efficiency of the untreated control was 3.62 x 10^−3^ with a standard deviation of 2.32 x 10^−3^. **B**. An ICE*Bs1* donor (CAL874) was mated with an ICE*Bs1*^0^ recipient in which the WTA biosynthesis operon *tagAB* had been placed under the control of Pspank (MMH584). Expression of *tagAB* was controlled by growing the recipient strain at the indicated IPTG concentrations. A wild type recipient (CAL89) was used as a control. Average mating efficiency with the wild type control was 6.07 x 10^−4^ with a standard deviation of 2.26 x 10^−4^. **C**. The Pspank-*tagAB* donor strain from Figure 2B (MMH578) was mated with the Pspank-*tagAB* recipient strain from 3B (MMH584). Both strains were grown at the indicated IPTG concentration. ND indicates relative mating efficiency was below the limit of detection (< 0.0007 and < 0.0008 for the 10 µM and 0 µM treatments, respectively). Average mating efficiency of the wild type control was 1.29 x 10^−3^ with a standard deviation of 8.19 x 10^−4^. **D**. The effect of inhibiting WTA biosynthesis on a Tn*916* donor (CMJ253) or recipient (MMH676) was tested by adding tunicamycin to growing cells 60 minutes prior to mating. Average mating efficiency of the untreated control was 4.49 x 10^−6^ with a standard deviation of 2.09 x 10^−6^.

We found that WTAs themselves are necessary for efficient ICE transfer, rather than WTA biosynthesis. We inhibited WTA biosynthesis without substantially depleting cells of WTAs by treating ICE*Bs1* donors with tunicamycin for one minute prior to mating. Under these conditions, there would be no substantial decrease in the amount of WTAs, but biosynthesis should be inhibited. There was little or no defect in conjugation of ICE*Bs1* under these conditions (Fig. 3A). Together, our results demonstrate that WTAs are required in donor cells for efficient transfer of ICE*Bs1*.

### WTAs are necessary in an ICE*Bs1* recipient for efficient transfer

We found that WTAs are also needed in a recipient for efficient acquisition of ICE*Bs1*. We treated recipients with tunicamycin for 1 hr and found that acquisition of ICE*Bs1* was reduced ∼50-fold (Fig. 3A). As with donors, blocking WTA biosynthesis in the recipient with tunicamycin treatment for only one minute prior to mating caused little or no defect in acquisition of ICE*Bs1* (Fig. 3A), indicating that WTAs themselves (and not simply WTA biosynthesis) are necessary in the recipient for efficient mating.

We also reduced WTAs in recipients by decreasing expression of *tagAB* (from a P*spank*-*tagAB* fusion) by growing cells in low concentrations (or in the absence) of IPTG for approximately six generations. With the lowest level of expression (no IPTG), the mating efficiency into the WTA-depleted recipients was decreased ∼200-fold relative to that of cells grown in 100 µM IPTG or wild type cells (Fig. 3B).

Reducing WTAs in both donors and recipients caused a defect in mating that was more severe than reduction in either donor or recipient alone (Fig. 3A, C). This was true following treatment of donors and recipients with tunicamycin (Fig. 3A) or growing both donors and recipients containing Pspank-*tagAB* in low concentrations of IPTG (Fig. 3C).

Based on these results, we conclude that WTAs in both donors and recipients are important for efficient conjugation of ICE*Bs1*. WTAs could be needed for the proper function, assembly, or regulation of the conjugation machinery, for establishing or maintaining cell-cell contact to enable mating, or for cell viability during mating.

### WTAs are not necessary for transfer of the broad host range ICE Tn*916*

We tested for effects of WTAs on conjugative transfer of Tn*916*, a small (∼18 kb) ICE that confers tetracycline resistance to its host. Activity of Tn*916* is increased several fold in the presence of tetracycline (Showsh and Andrews, 1992). Whereas the natural host of ICE*Bs1* appears to be limited to *B. subtilis*, Tn*916* is found in a broader range of gram-positive bacteria (Roberts and Mullany, 2009) and, although not its natural host, works quite well in *B. subtilis* (Christie *et al*., 1987; Celli and Trieu-Cuot, 1998; Johnson and Grossman, 2014).

We found that unlike ICE*Bs1*, Tn*916* transfer was not significantly affected when WTA biosynthesis was inhibited. We depleted WTAs in Tn*916* donors or recipients with tunicamycin (1 µg/ml) for 60 min. The mating efficiency following treatment of recipients was similar to that of untreated recipients, and that of donors appeared to increase slightly (Fig. 3D). This indicates that the decrease in transfer of ICE*Bs1* in response to WTA depletion is probably not due to a general effect on host physiology, or an inability of cells to contact each other, unless the presence of Tn*916* compensates for these alterations. This differential effect of WTAs on ICE*Bs1* compared to Tn*916* highlights a difference between conjugation mediated by these elements.

### An osmoprotective mating surface bypasses the need for WTAs in ICE*Bs1* conjugation

Cell wall hydrolases encoded by conjugative elements are essential for conjugation in gram-positive bacteria (Bantwal *et al*., 2012; Arends *et al*., 2013; Laverde Gomez *et al*., 2014; DeWitt and Grossman, 2014). Additionally, WTAs are important regulators of the activity of host-encoded cell wall hydrolases (autolysins) and are necessary for the proper localization of autolysins in several species (Yamamoto *et al*., 2008; Schlag *et al*., 2010; Frankel and Schneewind, 2012; Bonnet *et al*., 2018). Bacteria depleted of WTAs are more prone to autolysis and are more sensitive to treatment with lysozyme and autolysins (Bera *et al*., 2007; Atilano *et al*., 2010; Tiwari *et al*., 2018).

Based on the role of WTAs in modulating the activity of cell wall hydrolases, we hypothesized that WTA-depleted cells might have cell walls that are more sensitive to the formation of mating pairs. If true, then the decrease in conjugation efficiency should be suppressed (conjugation restored) under osmoprotective conditions that would enable cells to survive severe defects in their walls.

We found that the conjugation defect of WTA-depleted ICE*Bs1* donors was completely suppressed when matings were done on an osmoprotective surface (Fig. 4A). Matings were done on a standard mating surface (Spizizen’s salts, described in Experimental Procedures) or an osmoprotective mating surface that contained 20 mM MgCl2 and 0.5 M sucrose, buffered with 20 mM maleic acid pH 7 (MSM), an osmoprotective supplement that has been used to maintain protoplasts (lacking cell walls) and prevent bacterial cell death from osmotic stress (Wyrick and Rogers, 1973; Leaver *et al*., 2009). A deletion of a sucrose metabolism gene was incorporated into all strains used in these osmoprotection mating assays to prevent degradation of the sucrose osmoprotectant (Wolf *et al*., 2012). At the conclusion of the mating, cells were resuspended and diluted in MSM and then plated and grown on non-protective LB plates with the appropriate antibiotics to select for transconjugants.

**Figure 4.**
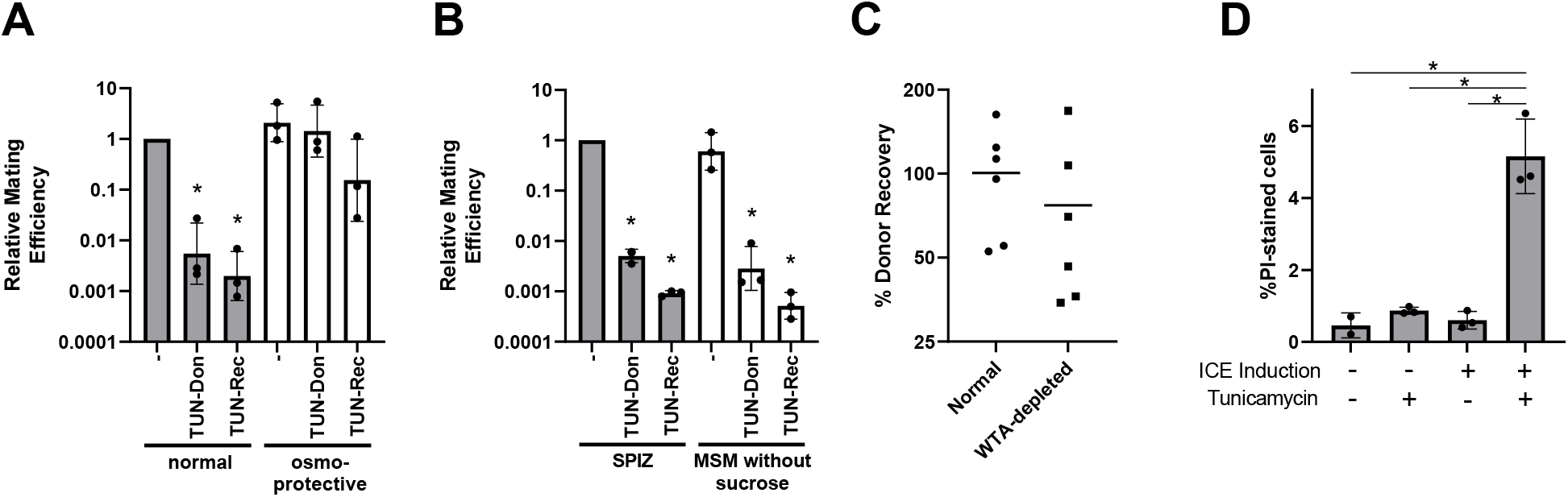
WTA-depleted ICE*Bs1* donors and recipients do not efficiently mate because they experience lethal cell envelope damage. **A**. The use of an osmoprotective mating surface eliminates the WTA-depletion ICE*Bs1* transfer defect. WTAs were depleted by treating ICE*Bs1* donors (MMH862) or recipients (MMH797) with 1 μg/ml tunicamycin for 60 minutes as in Figure 3A, and strains were mated on either a standard mating surface (1x Spizizen’s salts agar) or on an osmoprotective mating surface (1x MSM agar). Graph shows averages and standard deviations of log-transformed data from three biological replicates. * indicates average relative mating efficiency was significantly different from 1 (Student’s one-sample t-test, p < 0.05). Average mating efficiency of the wild type under non-protective conditions was 4.29 x 10^−3^ with a standard deviation of 2.14 x 10^−3^. **B**. WTA biosynthesis was inhibited by treating ICE*Bs1* donors (MMH862) or recipients (MMH797) with 1 μg/ml tunicamycin as in A. Strains were mated on either a standard mating surface (1x Spizizen’s salts agar) or on a 1x MSM agar surface lacking sucrose. Graph shows averages and standard deviations of log-transformed data from three biological replicates. * indicates average relative mating efficiency was significantly different from 1 (Student’s one-sample t-test, p < 0.05). Average mating efficiency for the untreated control (on SPIZ) was 1.63 x 10^−3^ with a standard deviation of 1.48 x 10^−3^. **C**. WTA-depleted ICE*Bs1* donors do not exhibit an observable drop in viability at the population level during mating. An ICE*Bs1* donor strain (MMH862) was or was not treated with tunicamycin and induced to mate with an ICE^0^ recipient (MMH797) as in A and B, and the number of viable post-mating donors was compared to the number of pre-mating donors. Each dot represents data from a single independent experiment (n = 6). The central bars represent the average. **D**. WTA-depleted cells engaging in ICE*Bs1* mating are more likely to sustain lethal membrane damage. ICE*Bs1* Δ*yddJ* cells with (MMH794) or without (MMH788) an IPTG-inducible ICE induction allele were cultured with IPTG and +/-tunicamycin, concentrated on an agar pad containing propidium iodide, and tracked for two hours via fluorescence microscopy. The percentage of PI-stained cells was recorded. Data are the averages of 3 biological replicates. Error bars indicate standard deviation. * indicates the averages of the two treatments are significantly different (Student’s two-sample t-test, p < 0.05).

As described above, treatment of donors with tunicamycin (1 µg/ml) for sixty minutes to deplete WTAs caused a mating defect under standard mating conditions (Fig. 4A). In contrast, this defect was fully suppressed under osmoprotective conditions and mating efficiencies were indistinguishable from those of cells without tunicamycin treatment (Fig. 4A).

Similarly, we found that the mating defect associated with WTA-depleted recipients was largely suppressed when matings were done on an osmoprotective mating surface (Fig. 4A). There was still a ∼6-fold drop in conjugation efficiency with WTA-depleted recipients on the osmoprotective mating surface relative to untreated cells (Fig. 4A). This drop was likely due to death of transconjugants that did not sufficiently recover from cell wall damage before the shift to non-protective conditions (LB agar).

Suppression of the mating defect on the osmoprotective surface (MSM) was due to osmoprotection by sucrose and not an effect of MgCl2 or the malate buffer. When sucrose was omitted from the mating surface, there was no rescue of the conjugation defect caused by depletion of WTAs (Fig. 4B). These results support the model that WTA-depleted cells die from osmotic stress. Furthermore, we conclude that WTA-depleted donors have significantly decreased ability to transfer ICE*Bs1*, perhaps because they die before they can successfully participate in conjugation. This could be due to overall death of all or the vast majority of ICE*Bs1-*containing cells, or selective death of a subpopulation, perhaps those that form mating pairs with recipients.

### WTA-depleted cells in ICE*Bs1* mating pairs are more likely to die

We found that there was not a large decrease in the number of viable WTA-depleted donors during mating (non-protective). As above, ICE*Bs1* was activated and cells were simultaneously treated with tunicamycin. This tunicamycin treatment caused a mild (3.7-fold) decrease in CFUs relative to untreated cells (1.1 x 10^8^ CFU/ml for untreated cells versus 3.0 x 10^7^ CFU/ml). We suspect that this drop in CFUs following tunicamycin treatment was due, at least in part, to defects in cell separation after division. We combined these donors with an ICE^0^ recipient on a (non-protective) mating surface. The percentage of viable donors recovered after mating was similar between cells with and without tunicamycin treatment (Fig. 4C), although there appeared to be a small drop in recovery of the tunicamycin-treated cells. Based on these results, we conclude that there is not a large drop in viability of the population of donors.

This population-level observation from the conclusion of a mating protocol does not reflect what occurs at the single-cell level. Although the vast majority of ICE*Bs1*-containing cells are potential donors, only a small number (∼1%) successfully participate in conjugation under conditions used here. Based on the results above, we hypothesized that WTA-depleted donors that are part of a mating pair likely undergo cell death. This would represent death of a small fraction of the population that would not be readily observed by bulk population-based viability assays.

We used propidium iodide (PI) staining and fluorescence microscopy to monitor death (loss of cell envelope integrity) of single cells. We induced a population of ICE*Bs1* donors to mate, concentrated them at high density on an agar pad containing propidium iodide, and monitored the cells for two hours to track the number of envelope-damaged (PI-stained) cells. Because mating is ordinarily a rare event, we used a monoculture of ICE*Bs1* donors lacking the ICE gene *yddJ*, which encodes a protein that would normally block the ICE+ cell from serving as a recipient in a mating pair (Avello *et al*., 2019). This allowed all cells to potentially serve as donors and recipients, thereby significantly increasing the frequency of conjugation. When ICE*Bs1* was induced in WTA-depleted cells under these conditions, we observed an approximately 8-fold increase in the incidence of PI-stained bacteria, indicating that WTA-depleted cells are more likely to die under conditions that support ICE*Bs1* transfer (Fig. 4D).

### The activity of the ICE*Bs1* conjugation machinery is sufficient to kill WTA-depleted donors and recipients

Our results indicate that WTA-depleted donors and recipients are defective in ICE*Bs1* mating due to envelope damage. In the case of matings using WTA-depleted recipients, this could be because 1) WTA-depleted recipients never acquire ICE*Bs1*, likely because forming a mating pair is lethal to WTA-depleted cells, or because 2) WTA-depleted recipients acquire ICE*Bs1* and become transconjugants but subsequently die, perhaps due to the expression of ICE*Bs1* genes or the transconjugant becoming a new donor.

We found that transfer of ICE*Bs1* into WTA-depleted recipients was not required for the decrease in conjugation efficiency. We measured mobilization of the plasmid pC194 by the ICE*Bs1* conjugation machinery into WTA-depleted recipients. In these experiments, we used a mutant ICE*Bs1* that is unable to excise from the chromosome (Δ*attR*) and that lacks a functional origin of transfer (Δ*oriT*). When activated, this ICE mutant still expresses the conjugation machinery and is able to mobilize several plasmids that do not encode their own conjugation system, including pC194 (Lee *et al*., 2012). We activated ICE*Bs1* gene expression and measured mobilization of pC194 into recipient cells. There was a ∼100-fold decrease in mobilization efficiency of pC194 into WTA-depleted (tunicamycin-treated) recipients compared to untreated recipients (Fig. 5). Based on these results, we conclude that the decrease in conjugation efficiency into WTA-depleted recipients is not due to transfer of ICE*Bs1* into recipients and subsequent death of the new transconjugant; rather, the defect is likely due to death of transconjugants caused by the formation of mating pairs or the act of transferring any DNA.

**Figure 5.**
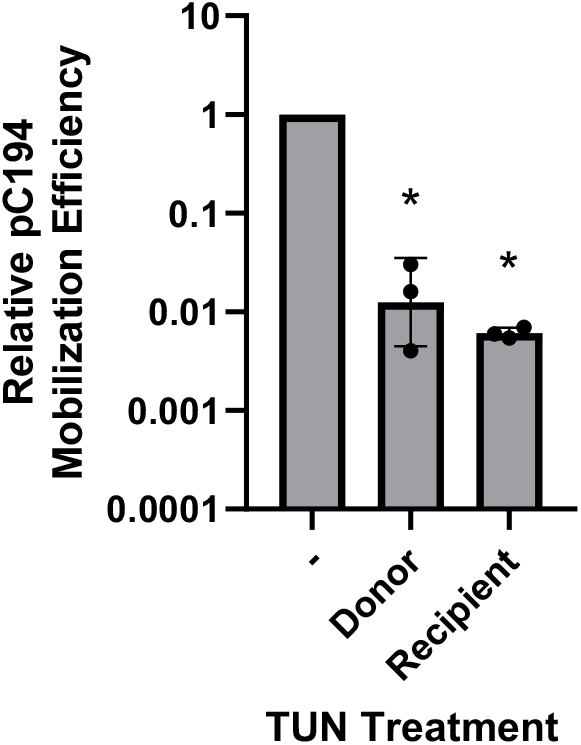
Mobilization of the non-conjugative plasmid pC194 by a locked-in ICE*Bs1* is negatively affected by WTA depletion. A strain containing both the mobilizable plasmid pC194 and a locked-in version of ICE*Bs1* (MMH868) was mated with an ICE*Bs1*^0^ recipient strain (MMH676). WTAs were depleted in either the donor or recipient strain via treatment with 1 μg/ml tunicamycin for 60 minutes as in Figure 3A. Relative mobilization efficiency was calculated as the number of transconjugants (Cm^R^ Strep^R^ CFUs) per initial donor relative to an untreated control. Graphs shows averages and standard deviations of log-transformed data from three biological replicates. * indicates relative mobilization efficiency was significantly different from 1 (Student’s one-sample t-test, p < 0.05). Average mobilization efficiency of the untreated control was 1.39 x 10^−3^ with a standard deviation of 6.21 x 10^−4^.

We performed similar experiments with WTA-depleted donor cells in which ICE*Bs1* could not transfer, but the ICE-encoded conjugation machinery could mobilize pC194. Again, there was an ∼100-fold decrease in mobilization efficiency as measured by acquisition of pC194 (Fig. 5). These results indicate that WTA-depleted donors are defective in transfer. Together with the results above, we infer that this defect is due to donor cell death in mating pairs.

## Discussion

We used a CRISPRi screen to identify essential genes in *B. subtilis* that are needed for efficient conjugation of ICE*Bs1*. We found that WTAs are necessary in both donor and recipient cells for efficient transfer of ICE*Bs1*. Under mating conditions, likely in mating pairs, there was significant death of WTA-depleted cells. Cell death and the need for WTAs for conjugation were reduced or eliminated when matings were done in osmoprotective conditions. Together, our results indicate that WTA-depleted cells fail to mate because they die from damage to their cell envelope.

The defect in conjugation of WTA-depleted cells was dependent on the ICE*Bs1* conjugation machinery, but did not depend on ICE*Bs1* DNA being transferred. We found that WTA-depleted cells were defective in transfer of a plasmid that normally can be transferred by the ICE*Bs1* conjugation machinery. In contrast, there was little or no defect in transfer of Tn*916*. These results indicate that the primary effect of WTAs is likely not to enable cell-cell contact (unless Tn*916* has some unknown function that does this). Rather, WTAs are important for proper function or control of the conjugation machinery, and that there is something fundamentally different about the activity of the ICE*Bs1* conjugation machinery compared to that of Tn*916* (see below).

### Possible mechanisms of conjugation-dependent toxicity in WTA-depleted cells

WTAs are important regulators of cell wall hydrolases across bacterial species, and WTA-depleted gram-positive bacteria have previously been demonstrated to be more sensitive to autolysin and lysozyme treatment (Brown *et al*., 2013). WTAs are also important for the proper localization of autolysins. Some *B. subtilis* cell wall hydrolases appear to be excluded from binding peptidoglycan decorated by WTAs, and the enzymes are mislocalized in the absence of WTAs (Yamamoto *et al*., 2008; Kasahara *et al*., 2016). It is possible that WTAs similarly regulate the activity of the bifunctional cell wall hydrolase CwlT that is encoded by ICE*Bs1*, and that mis-regulation of CwlT, perhaps in combination with mis-regulation of other cell wall hydrolases in WTA-depleted cells, causes the lethal envelope damage during conjugation. It is challenging to test the role of CwlT on conjugation efficiencies of WTA-depleted cells because *cwlT* is required for conjugation (DeWitt and Grossman, 2014).

WTAs also have a role in cell wall biosynthesis, and WTA-depleted *B. subtilis* cells exhibit irregularities in cell wall thickness and severe cell shape defects (D’Elia *et al*., 2006). It is possible that WTA-depletion in *B. subtilis* results in an unusually fragile cell wall, and that WTA-depleted *B. subtilis* is consequently much more sensitive to the normal cell wall modification process that occurs during conjugation.

### Activity of the conjugation machinery

Our results have interesting implications concerning the activity of the conjugation machinery and its effects on donor and recipient cells. First, cell death of WTA-depleted donors occurs only during mating conditions, and in a small subpopulation of cells. This indicates that the conjugation machinery is only activated in the presence of recipients, likely only in mating pairs. This is consistent with analyses of other elements that indicate the conjugation machinery is activated by cell-cell contact [reviewed in: (Christie *et al*., 2014)].

Additionally, if the decrease in acquisition of ICE*Bs1* by WTA-depleted recipients is due to killing that is mediated by the ICE*Bs1-*encoded cell wall hydrolase CwlT, then this indicates that the donor-encoded hydrolase is acting on recipient cells. There are other interpretations of this result: for example, the WTA-depleted cell’s own autolysins could be hyper-active, and active conjugation machinery from the donor could exacerbate the effects of hyper-active autolysins in recipients in mating pairs, thereby causing cell death.

Although CwlT appears to be the most likely cause of the conjugation-dependent toxicity among WTA-depleted cells, there are alternative explanations which cannot be ruled out. For example, WTA-depletion in *B. subtilis* sensitizes the cells to PBP-targeting antibiotic methicillin (Farha *et al*., 2013), and it is possible that the ICE*Bs1* conjugation machinery comparably interferes with cell wall biosynthesis in a way that is incompatible with WTA depletion. Alternatively, ICE*Bs1* conjugation could involve inactivating or modifying teichoic acids in a way that is lethal to WTA-depleted bacteria.

### Different ICEs respond differently to WTA depletion

We found that Tn*916* transfer was not negatively impacted by WTA-depletion, in contrast to our results with ICE*Bs1*. An important difference between the two elements that might contribute to this observation is that transfer of Tn*916* is less efficient than that of ICE*Bs1*. It is possible that less conjugation machinery is made when Tn*916* becomes transcriptionally active, and therefore the formation of mating pairs is not as stressful with the Tn*916*-encoded conjugation machinery as it is with that from ICE*Bs1*. It is also possible that the relevant components of the Tn*916* conjugation machinery are not regulated by *B. subtilis* WTAs, perhaps reflective of the broad host range of Tn*916*. It remains to be determined what it is about the ICE*Bs1* and Tn*916*-encoded conjugation machineries that make them respond so differently to *B. subtilis* WTAs.

Recent studies of ICE*St3* of *Streptococcus thermophilus* found that deleting a *tagO*-like gene, which might result in a decrease in WTAs, resulted in complex effects on transfer efficiency. Deletion in donors caused a decrease, but deletion in recipients caused an increase in conjugation efficiency (Dahmane *et al*., 2018). It is not clear if these effects are alleviated by osmoprotective conditions and if they are related to the effects described here for ICE*Bs1*.

### Conjugation and cell envelope stress

Connections between conjugation and envelope stress have long been known. One classic example is in *E. coli* where excessive transfer of F plasmids into F-recipients can lead to death of the recipient, a phenomenon called lethal zygosis (Skurray and Reeves, 1973). Subsequent studies found that radiolabeled peptidoglycan components are released into the medium when lethal zygosis occurs, indicating that the mechanism of death is due to damage of the cell envelope (Ou, 1980). Furthermore, activation of the F-plasmid sensitizes cells to certain envelope-disrupting antimicrobials (i.e. bile salts) (Bidlack and Silverman, 2004). The F plasmid has also been demonstrated to encode the means to upregulate the cell envelope stress response pathway in the host bacterium, which likely helps protect host cells from envelope stress caused by the F-encoded conjugation machinery (Grace *et al*., 2015).

Our results indicate that WTAs have an important role in protecting *B. subtilis* against envelope stress caused by the ICE*Bs1* conjugation machinery. We suspect that WTAs in other gram-positive bacteria also have a role in conjugation, and that role very likely will depend on aspects of the specific conjugation machinery assembled in the cell envelope.

## Methods

### Media and growth conditions

*B. subtilis* strains were grown at 37°C with shaking in LB medium. Experimental cultures were started from 3 ml LB exponential phase cultures inoculated from a single colony.

Where needed, *B. subtilis* strains were grown in LB at the following antibiotic concentrations for selection or maintenance of marked alleles: kanamycin (5 μg ml^-1^), streptomycin (100 μg ml^-^ 1), spectinomycin (100 μg ml^-1^), chloramphenicol (5 μg ml^-1^), and a combination of erythromycin (0.5 μg ml^-1^l) and lincomycin (12.5 μg ml^-1^) to select for macrolide-lincosamide-streptogramin (MLS) resistance and erythromycin resistance.

Tunicamycin was used at 1 μg ml^-1^ to inhibit WTA synthesis.

The osmoprotective supplement MSM (0.5 M sucrose, 20 mM MgCl2, buffered with 20 mM maleic acid pH 7) (Wyrick and Rogers, 1973; Leaver *et al*., 2009) was used where indicated. It was added from a 2x stock.

### Strains and alleles

*Escherichia coli* strain AG1111 (MC1061 F’ *lacI*^q^ *lacZM15* Tn*10*) was used for routine cloning and plasmid construction.

The *B. subtilis* strains used in this study are listed in Table 1. Strains were constructed using natural transformation (Harwood and Cutting, 1990). All *B. subtilis* strains are derivates of JH642 and contain tryptophan and phenylalanine auxotrophies (*trpC2 pheA1*) (Brehm *et al*., 1973; Smith *et al*., 2014). Many of the alleles used in this study have been described previously and are briefly summarized below.

**Table 1.**
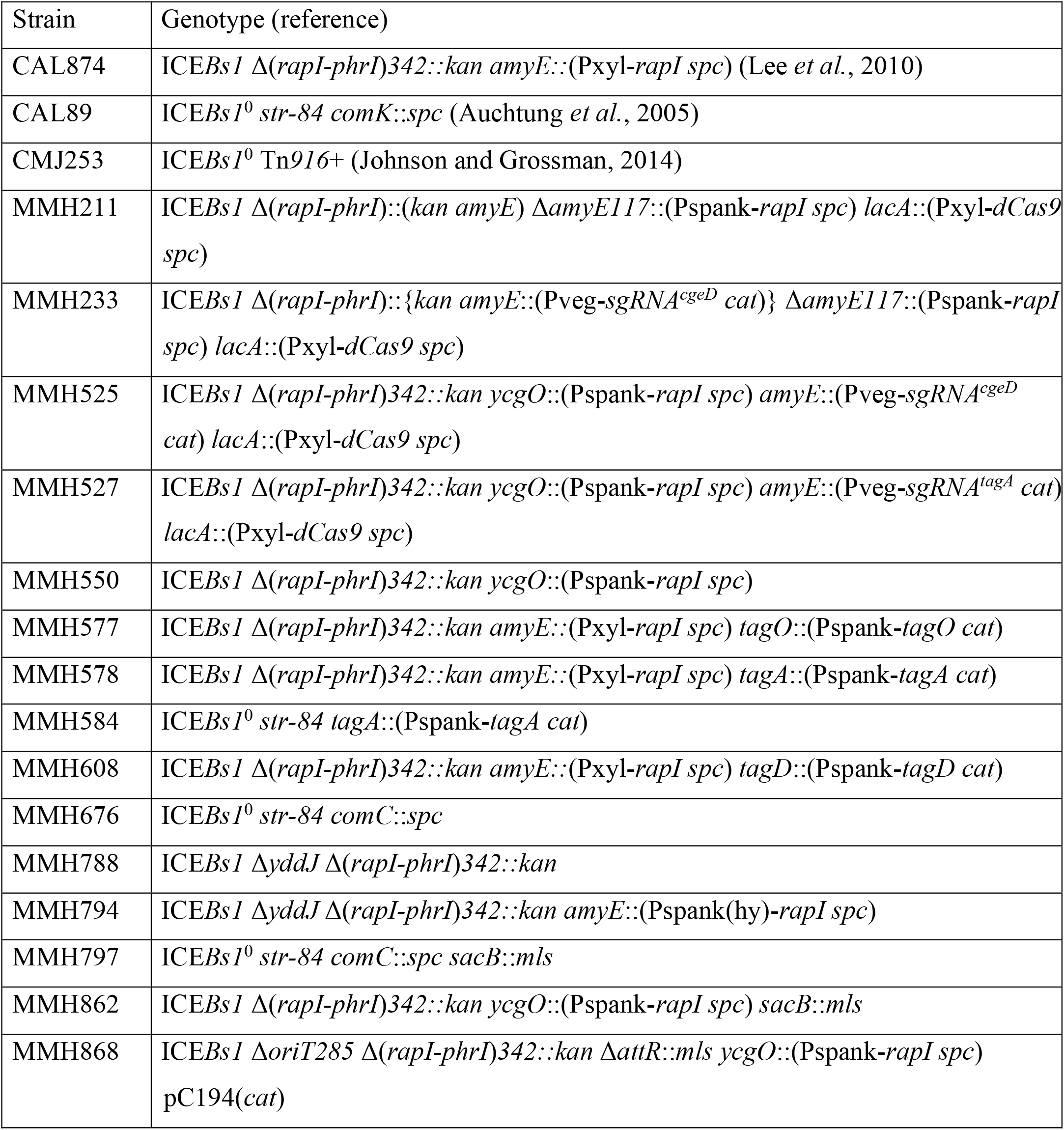
*B. subtilis* strains.

Donor strains used in standard ICE*Bs1* mating assays contained the allele Δ(*rapI-phrI*)*342*::*kan* (Auchtung *et al*., 2005). ICE*Bs1* was activated in donor strains by inducing expression of *rapI* from one of three promoter fusions: the LacI-repressible-IPTG-inducible promoters Pspank-*rapI* or Pspank(hy)-*rapI* (Auchtung *et al*., 2005), or the xylose-inducible promoter Pxyl-*rapI* (Berkmen *et al*., 2010). For the first two, *rapI* expression was induced by adding Isopropyl-β-D-thiogalactopyranoside (IPTG, Sigma) to a final concentration of 1 mM. Pxyl-*rapI* expression was induced via the addition of xylose to a final concentration of 1% w/v. ICE*Bs1*-containing strains used in live-cell microscopy also contained a deletion of *yddJ* (Avello *et al*., 2019).

Recipient strains were derived from the ICE*Bs1*-cured (ICE*Bs1*^0^) strain JMA222 (Auchtung *et al*., 2005) and were streptomycin resistant (*str84*) to facilitate counterselection during mating experiments. Recipient strains also contained *spc*-marked null alleles of competence genes *comK* or *comC* to prevent natural transformation.

ICE*Bs1* donor and recipient strains that were used in osmoprotective mating assays also contained a deletion-insertion of *sacB* (Δ*sacB*::*erm*) (Koo *et al*., 2017) to prevent degradation of sucrose (Wolf *et al*., 2012). A strain containing this allele was obtained from the Bacillus Genetic Stock Center (www.bgsc.org).

#### *ycgO*::{Pspank*-rapI* (*spc*)}

We constructed a *spc*-marked Pspank-*rapI* allele integrated into the nonessential *B. subtilis* gene *ycgO*. We used a previously described Pspank-*rapI* (*spc*) allele as a template for PCR amplification (Auchtung *et al*., 2005). The amplified allele was joined with *ycgO* flanking sequences by isothermal assembly (Gibson *et al*., 2009), and the construct was introduced to wild type *B. subtilis* via natural transformation selecting for spectinomycin resistance.

#### comC::spc

We constructed a *comC* deletion-insertion allele, extending from 324 bp upstream and 26 bp downstream of the *comC* open reading frame, with the *aad9* (*spc*) gene from pMagellan6. The allele was constructed by joining the appropriate *comC* flanking sequences with the amplified *aad9* gene by isothermal assembly, and was moved into wild type *B. subtilis* via natural transformation selecting for spectinomycin resistance.

#### Pspank*-tag* alleles

We constructed a set of three fusions that placed each of three endogenous WTA biosynthesis operons (*tagO, tagAB, tagDEF*) under the control of the IPTG-inducible promoter Pspank. In each case, Pspank was introduced upstream of the first gene in the operon by single cross-over integration. We used a plasmid (pJCL86; lab collection) that contains Pspank and *lacI* inserted into the backbone of pAG58 (Jaacks *et al*., 1989). A short region of the 5’UTR encompassing the predicted ribosome binding site and a few hundred bp of the 5’ region of the ORFs of *tagO, tagA*, and *tagD* were each amplified from wild type *B. subtilis* genomic DNA. The region amplified from each gene corresponded to the following: 19 bp upstream to 357 bp downstream of the *tagO* translation start site, 24 bp upstream to 277 bp downstream of the *tagA* translation start site, and 25 bp upstream to 157 bp downstream of the *tagD* translation start site. Each segment was inserted between the SphI and HindIII sites of pJCL86 via isothermal assembly, yielding plasmids pMMH558 (*tagO*), pMMH559 (*tagA*), and pMMH605 (*tagD*). The plasmids were transformed into wild type *B. subtilis* selecting for chloramphenicol resistance in the presence of 1 mM IPTG. Proper integration of each plasmid was confirmed by diagnostic PCR and DNA (Sanger) sequencing. The resulting *B. subtilis* strains grew normally in the presence of 1 mM IPTG, and exhibited severe growth defects in the absence of IPTG. Analysis of these strains by light microscopy following a transition out of IPTG-containing growth medium confirmed that all three strains exhibited the distinctive cell shape and separation defects characteristic of *B. subtilis* cells depleted of WTAs (D’Elia *et al*., 2006).

#### Construction of strains for mobilization of pC194

We constructed a *B. subtilis* strain containing the plasmid pC194 and a mutant of ICE*Bs1* that is unable to excise (Δ*attR*, ‘locked-in’) and without a functional origin of transfer (Δ*oriT*). This strain was made by moving the *ycgO*::{Pspank-*rapI* (*spc*)} allele into the ICE*Bs1*^0^ strain JMA222, creating strain MMH863. A version of ICE*Bs1* containing three mutations, Δ(*rapI-phrI*)*342*::*kan*, Δ*oriT*, and Δ*attR*::*mls*, was moved into MMH863 via natural transformation, and pC194 was subsequently introduced via natural transformation. The unmarked *oriT* deletion in this element has been described (Jones *et al*., 2020). The Δ*attR*::*mls* allele was constructed via isothermal assembly using *mls* from pCAL215 (Lee *et al*., 2007) as a template, and has the same deletion boundaries as a previously reported Δ*attR*::*tet* allele (Lee and Grossman, 2007).

#### Construction of CRISPRi ICE*Bs1* donor library

The Pxyl-*dcas9* and Pveg-*sgRNA* alleles used to generate the CRISPRi knockdown library of ICE*Bs1* donor strains were previously described and a generous gift from Peters *et al*., (Peters *et al*., 2016). The initial library contains a set of 299 plasmids, each containing a Pveg-*sgRNA* allele with a unique 20 bp targeting region, a *cat* marker conferring chloramphenicol resistance, and the appropriate flanking homology needed to integrate the sgRNA allele into the *B. subtilis* chromosome at *amyE* via double crossover. We utilized a pooled collection of these plasmids to generate our library of Pveg-*sgRNA* alleles integrated into ICE*Bs1*.

Our strategy was to insert *amyE* into ICE*Bs1*, delete *amyE* from the chromosome, and then recombine the Pveg-*sgRNA* library into *amyE* in ICE*Bs1*. We constructed a deletion-insertion of *amyE* (Δ*amyE117*::{Pspank-*rapI* (*spc*)}) that replaces the entire chromosomal gene and flanking noncoding regions with Pspank-*rapI*. This allele was constructed using a previously described Pspank-*rapI* (*spc*) allele (Auchtung *et al*., 2005) as a template for PCR amplification, and by joining the amplified product to *amyE* flanking homology via isothermal assembly. The construct was transformed into *B. subtilis* selecting for resistance to spectinomycin. The allele was confirmed by sequencing PCR-amplified DNA and verified functionally to activate ICE*Bs1* following addition of IPTG.

We used isothermal assembly to construct a *kan*-marked copy of *amyE* with flanking sequences needed to integrate the allele into *rapI-phrI* of ICE*Bs1*. This allele (Δ(*rapI-phrI*)::{*amyE kan*}) was designed such that the deletion boundaries and orientation of the *kan* cassette would be identical to Δ(*rapI-phrI*)*342*::*kan*.

The pooled library of Pveg-*sgRNA* plasmids was linearized with KpnI-HF (NEB), and was incorporated into the ICE::*amyE* locus by transformation into strain MMH211 and selecting for resistance to chloramphenicol. We recovered ≥ 1.5 x 10^5^ transformants in total. Transformants were resuspended and pooled in MOPS-buffered S750 defined minimal medium (Jaacks *et al*., 1989) lacking a carbon source (1x S750 + metals) and frozen in aliquots for future use.

We constructed one Pveg-*sgRNA* allele for use as a control, with a 20 bp targeting region corresponding to the nonessential *B. subtilis* gene *cgeD*. To construct this allele we used inverse PCR as previously described (Larson *et al*., 2013) using pJMP2 as a template for amplification (Peters *et al*., 2016), creating plasmid pMMH221. The plasmid was linearized with KpnI-HF and transformed into the appropriate *B. subtilis* strains via selection with chloramphenicol. A copy of ICE*Bs1* with the Pveg-*sgRNA*^cgeD^ allele incorporated into the ICE::*amyE* integration site was confirmed to transfer normally.

### CRISPRi library mating

A lawn of the CRISPRi ICE*Bs1* donor library was started from freezer stocks on the day before the experiment and grown overnight at room temperature on an LB plate. The following day, the lawn was used to start a culture at OD600 = 0.02 in LB supplemented with 0.01% xylose to stimulate transcription of Pxyl-*dcas9* and with kanamycin to ensure maintenance of the *kan*-marked ICE*Bs1*. When cultures reached an OD600 of 0.15, ICE*Bs1* was activated by addition of IPTG (1 mM) for 1 hr to induce expression of Pspank-*rapI*.

Mating of the library into the streptomycin resistant ICE-cured recipient strain CAL89 was conducted using a standard mating procedure (see below). An aliquot of the donor culture was harvested for analysis immediately prior to mixing donors and recipients. At the conclusion of the mating, the cells on the filter were resuspended and plated on LB agar containing kanamycin and streptomycin to select for transconjugants and incubated overnight at 37°C. Transconjugants were then resuspended in 1x S750 + metals and pooled, and an aliquot of the resuspension was harvested for sequence analysis.

### Amplification and sequencing of pooled sgRNA alleles

We used Qiagen DNeasy Blood and Tissue kits to extract DNA from the pre- and post-mating samples, following the manufacturer instructions for the use of the kit with gram-positive bacteria. We used KAPA HiFi MasterMix to amplify the *sgRNA* alleles from the DNA samples, adhering to the manufacturer protocol. We used 300 ng of sample DNA as a template and ran the reactions for 20 PCR cycles. The primers used to amplify the sgRNA alleles were oMH238 (5’-AATGATACGGCGACCACCGAGATCTACACGGGCGGGAATGGGCTCGTGTTGTACAATAAATGT-3’) and oMH239 (5’-CAAGCAGAAGACGGCATACGAGATXXXXXXGCCAGCCGATCCTCTTCTGAGATGAG TTTTTGTTCG-3’). The 5’ ends of these oligos encode the Illumina adaptor sequences, and the X’s correspond to multiplexing barcodes. The resulting amplicons were purified with SureSelect AMPure beads according to manufacturer instructions. The sgRNA amplicons were sequenced with an Illumina MiSeq, using oMH240 to sequence the variable region of the sgRNA allele (5’-GGGCGGGAATGGGCTCGTGTTGTACAATAAATGT-3’) and oMH241 to sequence the multiplexing barcode (5’-CGAACAAAAACTCATCTCAGAAGAGGATCGGCTGGC-3’).

### Mating assays

Mating assays were conducted as previously described (Auchtung *et al*., 2005; DeWitt and Grossman, 2014; Johnson and Grossman, 2014), with minor modifications. Experimental cultures of donor and recipient strains were started in LB at an OD600 of 0.02 and grown with shaking at 37°C. Kanamycin was added to cultures of strains containing a *kan*-marked ICE*Bs1* to ensure maintenance of the element. ICE*Bs1* activation was induced at an OD600 of 0.15 in donor cultures by adding either 1mM IPTG or 1% xylose. One hour after induction, 2.5 OD600 equivalents of donors were combined with an equal amount of recipients. The mixture of donors and recipients was collected on an nitrocellulose filter via vacuum filtration, and washed with 5 ml 1x Spizizen’s salts (2 g/l (NH4)SO4, 14 g/l K2HPO4, 6 g/l KH2PO4, 1 g/l Na3 citrate-2H2O, and 0.2 g/l MgSO4-7H2O) (Harwood and Cutting, 1990). The filter was placed on a 1x Spizizen’s salts 1.5% agar plate (without a carbon source) and incubated at 37°C for 2 h. The cells were resuspended in 5 ml 1x Spizizen’s salts. Transconjugants were quantified by serially diluting the resuspension in 1x Spizizen’s salts, spreading the cells on an LB 1.5% agar plate containing kanamycin and streptomycin (for ICE*Bs1* matings), and incubating the plates overnight at 37°C. Donors were quantified immediately prior to combining donor and recipient cells: an aliquot of the donor culture was serially diluted and plated on an LB 1.5% agar plate containing kanamycin and grown overnight at 37°C. Mating efficiency was calculated as the number of transconjugants (kanamycin- and streptomycin-resistant post-mating CFUs/ml) per initial donor (pre-mating donor CFUs/ml). Data reported are normalized to the mating efficiency of a wild type control strain assayed on the same day.

For mating assays done with individual strains with inducible CRISPRi-mediated knockdowns, Pxyl-*dcas9* expression was stimulated at the time of culture inoculation by supplementing the experimental cultures with 0.01% or 0.1% xylose.

For mating assays with strains in which the endogenous *tag* operons had been placed under the control of the IPTG-inducible promoter Pspank, the LB starter cultures were grown in the presence of 100 μM IPTG. The starter cultures were pelleted by centrifugation and washed twice with plain LB (no IPTG). Cells were resuspended in LB containing the indicated concentration of IPTG (ranging from 0 to 100 μM), and were used to inoculate LB cultures containing IPTG at the same concentration. LB agar plates were supplemented with 1 mM IPTG to obtain CFUs for strains with an IPTG-inducible promoter fused to an essential gene.

For mating assays with tunicamycin-treated strains, tunicamycin was added concurrently with ICE*Bs1* induction to a final concentration of 1 μg ml^-1^.

For mating assays with Tn*916*, ICE activation was stimulated with 2.5 μg ml^-1^ tetracycline, and tetracycline was used in the selection for donors and transconjugants (instead of kanamycin).

For mating assays done on an osmoprotective mating surface, the 1x Spizizen’s salts 1.5% agar plate was substituted for a 1x MSM 1.5% agar plate, and the cells were resuspended and diluted post-mating in 1x MSM instead of 1x Spizizen’s salts. Pre- and post-mating CFUs were grown out by plating on non-osmoprotective LB 1.5% agar plates as described above. Sucrose was omitted from the 1x MSM solid and liquid media for the relevant experimental controls.

### pC194 mobilization assays

pC194 mobilization assays were carried out in essentially the same manner as ICE*Bs1* mating assays. Donor cultures were grown with chloramphenicol instead of kanamycin to maintain pC194 (containing *cat*). Donors and transconjugants were measured by plating on appropriate selective media.

### Live cell microscopy

Live-cell microscopy was done largely as previously described (Babic *et al*., 2011) with minor modifications. Cultures of ICE*Bs1*+ Δ*yddJ* strains were grown in LB as described above. Strains either did or did not contain *amyE*::Pspank(hy)-*rapI*. All strains were treated with 1 mM IPTG for 1 hr at an OD600 of 0.15. For WTA-depletion, tunicamycin (1 μg/ml) was added concurrently with IPTG. After 1 hr, cells were pelleted in a tabletop centrifuge at 14000 rpm, washed once in 1x S750 + metals, and resuspended in 50 μl 1x S750 + metals. 1 μl of the resuspension was applied to an agar pad. The agar pad comprised 1.5% Noble Agar (Difco) dissolved in carbonless 1.5% 1x S750 + metals medium and contained 30 μM propidium iodide.

The agar pad was placed on a glass coverslip (VWR) such that the cells would be in contact with the coverslip, and the coverslip was attached to a microscope slide via a frame-seal slide chamber (Bio-Rad). The cells were observed via a Nikon Ti-E inverted microscope and using a CoolSnap HQ camera (Photometrics). Propidium iodide fluorescence was generated with a Nikon Intensilight mercury illuminator through an excitation and emission filter (Chroma; filter set 49008) The cells were monitored for PI-staining at 37°C for 2 h at 15 minute timepoints. At least 1,000 cells were monitored for each biological replicate.

## Acknowledgements

We thank Jason Peters and Carol Gross for their generous gift of the *B. subtilis* essential gene knockdown library which enabled this work, and Michael Laub, Gene-Wei Li, David Popham, and Mary Anderson for comments on the manuscript.

The research reported here was supported by the National Institute of General Medical Sciences of the National Institutes of Health under award number R01 GM050895 and R35 GM122538 to ADG. MMH was also supported, in part, by the NIGMS predoctoral training grant T32 GM007287. Any opinions, findings, and conclusions or recommendations expressed in this report are those of the authors and do not necessarily reflect the views of the National Institutes of Health.

