## Supplemental Table 1 for "A CRISPR interference screen reveals a role for cell wall teichoic acids in conjugation in *Bacillus subtilis*"

**Table S1. Results of CRISPRi screen.** The CRISPRi screen for essential host gene knockdowns that cause an acute ICE*Bs1* defect was carried out as described in the text and in Figure 1. Relative ability of a knockdown strain to donate ICE*Bs1* was monitored by using next generation sequencing to determine the relative abundance of an sgRNA in the transconjugant population versus the abundance of the sgRNA in the population of pre-mating donors. Gene knockdowns that caused an impactful ICE*Bs1* transfer defect (>4-fold change in sgRNA abundance) and that were not depleted from the donor population prior to mating (>0.01% of pre-mating donor population) are reported here. The proportion of the starting population that each sgRNA constitutes is also listed, as well as the *Subti*wiki functional annotation for each gene (http://subtiwiki.uni-goettingen.de/v3/) (Zhu and Stülke, 2018).

| Gene Name | Fold-Change After Mating (TC abundance / donor abundance) | Proportion of Pre-Mating Donor Population | Gene Function (Subtiwiki) |
|---|---|---|---|
| *tagA* | eliminated | $1.65 \times 10^{-3}$ | biosynthesis of teichoic acid |
| *rplL* | eliminated | $1.68 \times 10^{-4}$ | translation |
| *fusA* | eliminated | $1.74 \times 10^{-4}$ | translation |
| *rplF* | eliminated | $1.61 \times 10^{-4}$ | translation |
| *secE* | $7.73 \times 10^{-4}$ | $1.20 \times 10^{-3}$ | protein secretion |
| *mntA* | $1.12 \times 10^{-3}$ | $3.32 \times 10^{-3}$ | manganese uptake |
| *rpsC* | $1.66 \times 10^{-3}$ | $2.80 \times 10^{-4}$ | translation |
| *cysS* | $5.47 \times 10^{-3}$ | $4.26 \times 10^{-4}$ | translation |
| *groEL* | $8.04 \times 10^{-3}$ | $3.42 \times 10^{-3}$ | protein folding and re-folding |
| *infC* | 0.0100 | $1.86 \times 10^{-4}$ | translation |
| *groES* | 0.0105 | $3.60 \times 10^{-3}$ | protein folding and re-folding |
| *rplX* | 0.0194 | $4.56 \times 10^{-4}$ | translation |

| | | | |
|---|---|---|---|
| *rny* | 0.0531 | $2.25 \times 10^{-3}$ | RNA processing and degradation |
| *rplN* | 0.0566 | $2.06 \times 10^{-4}$ | translation |
| *ydiO* | 0.0588 | $3.14 \times 10^{-3}$ | BsuM DNA modification |
| *tufA* | 0.0748 | $1.62 \times 10^{-4}$ | translation |
| *ydiP* | 0.0822 | $2.85 \times 10^{-3}$ | BsuM DNA modification |
| *rplJ* | 0.0836 | $1.41 \times 10^{-3}$ | translation |
| *tagD* | 0.103 | $3.82 \times 10^{-3}$ | biosynthesis of teichoic acid |
| *gpsA* | 0.104 | $2.07 \times 10^{-4}$ | biosynthesis of phospholipids |
| *nusA* | 0.106 | $3.19 \times 10^{-3}$ | transcription |
| *secA* | 0.110 | $1.96 \times 10^{-3}$ | protein secretion |
| *tsf* | 0.115 | $4.46 \times 10^{-4}$ | translation |
| *glmS* | 0.117 | $3.50 \times 10^{-3}$ | cell wall synthesis |
| *murAA* | 0.117 | $8.72 \times 10^{-4}$ | peptidoglycan precursor biosynthesis |
| *rpsNA* | 0.120 | $8.90 \times 10^{-4}$ | translation |
| *nrdI* | 0.120 | $1.41 \times 10^{-3}$ | synthesis of deoxyribonucleotide triphosphates |
| *mreC* | 0.141 | $1.18 \times 10^{-3}$ | cell shape determination |
| *fabF* | 0.142 | $2.48 \times 10^{-3}$ | fatty acid biosynthesis |
| *murB* | 0.143 | $2.71 \times 10^{-4}$ | peptidoglycan precursor biosynthesis |
| *nrdE* | 0.144 | $5.61 \times 10^{-4}$ | synthesis of deoxyribonucleotide triphosphates |
| *trnI-Asn* | 0.148 | $8.08 \times 10^{-3}$ | translation |
| *asnB* | 0.178 | $9.67 \times 10^{-4}$ | control of peptidoglycan hydrolysis |
| *rplS* | 0.201 | $5.13 \times 10^{-4}$ | translation |
| *ffh* | 0.203 | $3.27 \times 10^{-3}$ | presecretory protein translation |
| *adk* | 0.206 | $1.90 \times 10^{-4}$ | ADP formation |
| *rpsH* | 0.218 | $1.00 \times 10^{-3}$ | ribosomal protein |
| *yneF* | 0.218 | $1.11 \times 10^{-3}$ | membrane protein of unknown function |

| | | | |
|---|---|---|---|
| *tagF* | 0.230 | $2.74 \times 10^{-3}$ | biosynthesis of teichoic acid |
| *alaS* | 0.232 | $7.59 \times 10^{-4}$ | translation |
| *parE* | 0.239 | $4.93 \times 10^{-4}$ | chromosome segregation and compaction |

Zhu, B., and Stülke, J. (2018) *Subti*Wiki in 2018: from genes and proteins to functional network annotation of the model organism *Bacillus subtilis*. *Nucleic Acids Res* **46**: D743–D748.
